# Palmitate-activated IRE1-XBP1 signaling limits DNA damage and is associated with H2A.X-linked DNA damage response transcripts in triple-negative breast cancer cells

**DOI:** 10.64898/2026.09.25.754433

**Authors:** Kevin Y. Chen, Elaina Gouin, Ava Erickson, Yogit Goyal, Samantha Zaloudek, Sean Foster, Caleb Sandum, S. Patrick Walton, Christina Chan

**Affiliations:** Department of Chemical Engineering and Materials Science, Michigan State University, MI, USA; Department of Biochemistry and Molecular Biology, Michigan State University, MI, USA; Department Microbiology, Genetics, & Immunology, Michigan State University, MI, USA; Institute of Quantitative Health Science and Engineering, Michigan State University, MI, USA

## Abstract

Palmitate-associated lipotoxic stress activates the endoplasmic-reticulum stress sensor protein IRE1, but how IRE1 signaling influences genotoxic-stress responses in triple-negative breast cancer remains unclear. Here, we investigated IRE1 and XBP1 signaling in MDA-MB-231 cells exposed to palmitate and etoposide. Under combined lipotoxic and genotoxic stress, XBP1 depletion increased DNA-damage accumulation in IRE1 wild-type cells but not in IRE1-deficient cells, whereas IRE1 re-expression reduced DNA damage. Pharmacologic IRE1 RNase inhibition also increased etoposide-associated DNA damage, supporting a protective role for IRE1 RNase-dependent signaling. IRE1 status and XBP1 manipulation differentially affected H2A.X and γH2A.X abundance, indicating context-dependent regulation of chromatin-associated DNA-damage signaling. At the RNA level, H2A.X was enriched in IRE1 immunoprecipitates, and the H2A.X-linked DNA-damage-response transcripts HUWE1 and BRCA2 also associated with IRE1 under combined stress. These findings link palmitate-activated IRE1-XBP1 signaling to genome-protective responses during genotoxic stress and identify a network of IRE1-associated DNA-damage-response transcripts in triple-negative breast cancer cells. Direct IRE1-mediated cleavage of these transcripts remains to be established.

## Introduction

Genome integrity is essential for normal cellular function, and genomic instability is a hallmark of cancer that drives chromosomal rearrangements, oncogene activation, and loss of tumor suppressor function (ref. 1, 2). Cancer cells frequently exhibit dysregulated DNA-damage-response (DDR) signaling, which can promote survival after genotoxic stress and reduce chemosensitivity (ref. 3, 4). Cancers, including triple-negative breast cancer (TNBC), commonly display genomic instability and DDR defects, complicating the use of cytotoxic chemotherapy, which remains central to its treatment (ref. 5, 6). Obesity is associated with TNBC risk and progression through several systemic and tumor-microenvironmental mechanisms, including altered lipid availability (ref. 7). Saturated fatty acids such as palmitate (PA) can induce lipotoxic cellular stress and an endoplasmic-reticulum (ER) stress response (ref. 8). In Hep3B liver cancer and MDA-MB-231 breast cancer cells, PA-induced inositol-requiring enzyme 1α (IRE1)-X-box binding protein 1 (XBP1) signaling has been shown to promote migration (ref. 9). These findings support a rationale for examining whether PA-induced IRE1 signaling also alters DDR responses during genotoxic stress in TNBC.

DNA double-strand breaks (DSBs) are highly cytotoxic lesions that can arise following treatment with genotoxic chemotherapeutics, including etoposide (ref. 10, 11). Upon DSB formation, the histone H2A variant H2A.X is phosphorylated at Ser139, generating γH2A.X, which spreads through chromatin flanking the break site (ref. 12, 13). This chromatin modification contributes to DNA-damage signaling and coordinates repair at damaged chromatin, allowing γH2A.X to function as both a marker of DNA damage and a platform for the assembly of DNA-repair factors (ref. 15). Overall H2A.X levels shape cellular DDR capacity (ref. 15–17). Genome maintenance at the chromatin level also relies on additional factors that safeguard replication forks and coordinate repair, including E3 ubiquitin-protein ligase (HUWE1) and breast cancer type 2 susceptibility protein (BRCA2). HUWE1 is recruited to stalled forks and drives H2A.X ubiquitination to support signaling and fork recovery during replication stress (ref. 18), while BRCA2 stabilizes stalled forks and enables recombination-based repair of broken DNA (ref. 19, 20). Together, H2A.X, HUWE1, and BRCA2 coordinate complementary DDR functions involving damage signaling and the protection and recovery of stalled replication forks (ref. 18–20).

Increasing evidence links DNA-damage signaling to ER proteostasis, the cellular system that maintains protein folding and ER homeostasis (ref. 21). Perturbation of ER proteostasis activates the unfolded protein response (UPR), which is coordinated by the ER stress sensors IRE1, protein kinase R-like endoplasmic reticulum kinase, and activating transcription factor 6 (ref. 22). The UPR reduces protein-folding demand, increases chaperone and ER-associated degradation capacity, and can promote cell death when adaptive signaling fails to restore homeostasis (ref. 23). In cancer, however, persistent UPR signaling can be co-opted to support tumor-cell survival (ref. 24), metastasis (ref. 9), and resistance to therapy (ref. 25).

Among the three UPR sensors, IRE1 is uniquely suited to link ER stress to post-transcriptional gene regulation through its cytosolic kinase and RNase domains (ref. 26, 27). Activated IRE1 catalyzes unconventional splicing of XBP1 mRNA to produce XBP1s, a transcription factor that promotes adaptive proteostasis programs (ref. 28, 29). IRE1 can also regulate selected transcripts through regulated IRE1-dependent decay (RIDD), an RNase-domain activity distinct from XBP1 splicing (ref. 30, 31). Previous study indicates that genotoxic stress can engage IRE1-dependent RNA decay and alter DDR-associated transcript stability, even in settings without robust XBP1 mRNA splicing (ref. 32). Although PA can activate IRE1–XBP1 signaling in cancer cells and genotoxic stress can activate IRE1-dependent RNA decay, it is unknown whether IRE1 activity is associated with changes in H2A.X/γH2A.X abundance and the expression of selected DDR-related transcripts. Furthermore, it is unclear whether PA alters IRE1/XBP1-dependent responses to etoposide-induced DNA damage.

In this study, we investigated how PA-associated stress modulates IRE1/XBP1 signaling and the DDR after exposure to etoposide in MDA-MB-231 cells, an invasive TNBC line established from a metastatic pleural effusion (ref. 33, 34). We tested whether the presence of IRE1 and XBP1 abundance or splicing influence DNA-damage accumulation and H2A.X/γH2A.X responses. We also assessed whether H2A.X and the H2A.X-linked DDR transcripts HUWE1 and BRCA2 associate with IRE1. We show that IRE1/XBP1 signaling shapes the cellular response to combined PA and etoposide stress and is associated with changes in DNA-damage accumulation, H2A.X/γH2A.X dynamics, and select DDR-associated transcripts.

## Materials and methods

### Cell culture

MDA-MB-231 human triple-negative breast cancer cells were obtained from the American Type Culture Collection (ATCC; HTB-26, Manassas, VA, USA). Cells were maintained in glucose-free Dulbecco’s modified Eagle medium (DMEM; Gibco, Thermo Fisher Scientific, Waltham, MA, USA; cat. no. 11966025) supplemented with 10% fetal bovine serum (FBS; Dot Scientific, Burton, MI, USA) and D-glucose to a final concentration of 5.5 mM (Gibco Glucose Solution, Thermo Fisher Scientific; cat. no. A2494001). Cells were cultured at 37 °C in a humidified incubator with 5% CO₂. IRE1-deficient (IRE1^−/−^) MDA-MB-231 cells were generated by CRISPR/Cas9-mediated gene editing and validated as previously described (ref. 9). The same IRE1^−/−^ cell population was used in this study. IRE1 WT cells refer to parental MDA-MB-231 cells. Experiments were performed using cells between passages 8 to 25.

### Cell treatments and transfection

Cells were transfected with 10 nM ON-TARGETplus Human *XBP1* siRNA SMARTpool (Horizon Discovery; cat. no. L-009552-00-0005) or ON-TARGETplus Non-targeting Control Pool (Horizon Discovery; cat. no. D-001810-10-05) using Lipofectamine 3000 Transfection Reagent (Thermo Fisher Scientific; cat. no. L3000150) according to the manufacturer’s instructions. Cells were incubated for 48 h after siRNA transfection before chemical treatment.

For overexpression and re-expression experiments, MDA-MB-231 IRE1^−\−^ cells were transfected with 1,000 ng of empty vector or plasmids encoding human IRE1α/ERN1, XBP1s, or XBP1u using Lipofectamine 3000 according to the manufacturer’s instructions. Cells were incubated for 24 h after plasmid transfection before chemical treatment. Equal total amounts of plasmid DNA were used across conditions.

Sodium palmitate (PA; Sigma-Aldrich; cat. no. P9767) was used at a final concentration of 0.1 mM. For PA+ETP experiments, cells were pretreated with PA for 8 h and subsequently co-treated with PA and 10 µM etoposide (ETP; Sigma-Aldrich; cat. no. E1383) for 16 h. ETP-only controls received 10 µM ETP for 16 h in the absence of PA. For pharmacologic experiments, WT MDA-MB-231 cells were treated with 30 µM 4μ8C (Sigma-Aldrich; cat. no. SML0949) or volume-matched DMSO vehicle for 3 h before, and during, a subsequent 3-h exposure to 10 µM ETP. Detailed cell-treatment and transfection procedures are provided in the Supplementary Methods.

### Immunoblotting

Cells were lysed on ice for 15 min in RIPA Lysis and Extraction Buffer (Thermo Fisher Scientific, Waltham, MA, USA; cat. no. 89901) supplemented with 1× Halt Protease and Phosphatase Inhibitor Cocktail (Thermo Fisher Scientific, Waltham, MA, USA; cat. no. 78440). Lysates were clarified by centrifugation at 21,130×g (max) for 20 minutes at 4 °C, and protein concentrations were determined using the Pierce BCA Protein Assay Kit (Thermo Fisher Scientific, Waltham, MA, USA; cat. no. 23225) according to the manufacturer’s instructions. Equal amounts of protein (15 µg/lane) were separated on 4–15% or 8–16% Mini-PROTEAN TGX precast gels (Bio-Rad Laboratories, Hercules, CA, USA) and transferred to 0.2-µm PVDF membranes using the Trans-Blot Turbo Transfer System and Ready-to-Assemble Mini PVDF Transfer Kits (Bio-Rad Laboratories) according to the manufacturer’s instructions.

Membranes were blocked in EveryBlot Blocking Buffer (Bio-Rad Laboratories, Hercules, CA, USA) for 5 min at room temperature and incubated with primary antibodies diluted 1:1,000 in EveryBlot Blocking Buffer for 1 h at room temperature. For H2A.X and γH2A.X immunoblotting, membranes were blocked in 5% BSA in 1× TBST for 1 h at room temperature and incubated with primary antibodies diluted 1:1,000 in 5% BSA/TBST overnight for 16 h at 4 °C. Membranes were washed 5×5 minutes in TBST and incubated with HRP-conjugated secondary antibodies diluted 1:10,000 in the respective blocking buffer for 1 h at room temperature.

Chemiluminescent signals were developed using Clarity Western ECL Substrate (Bio-Rad Laboratories, Hercules, CA, USA; cat. no. 1705061) and acquired using a ChemiDoc MP Imaging System (Bio-Rad Laboratories). Band intensities were quantified using Image Lab software (version 6.1.0; Bio-Rad Laboratories) and normalized to vinculin or β-actin. Antibody details are provided in Supplementary Table S1.

### Analysis of mRNA levels by real-time quantitative PCR

Total RNA was isolated using the miRNeasy Tissue/Cells Advanced Mini Kit (QIAGEN, Hilden, Germany; cat. no. 217604) according to the manufacturer’s instructions. RNA concentration and purity were assessed by NanoDrop spectrophotometry (Thermo Fisher Scientific), and samples with A260/280 ratios of 1.8–2.1 were used for cDNA synthesis. cDNA was generated from 200 ng total RNA using the High-Capacity RNA-to-cDNA Kit (Applied Biosystems, Thermo Fisher Scientific; cat. no. 4387406) according to the manufacturer’s protocol. RT-qPCR was performed using PowerTrack SYBR Green Master Mix (Applied Biosystems, Thermo Fisher Scientific; cat. no. A46109) on a QuantStudio 5 Real-Time PCR System (Applied Biosystems, Thermo Fisher Scientific), following the manufacturer’s instructions. Gene-specific primers targeted H2A.X, HUWE1, BRCA2, BLOC1S1, SPARC, XBP1 and 18S rRNA. H2A.X, HUWE1, and BRCA2 primer pairs were the same as those used for CLIP-qRT-PCR. Primer sequences are listed in Supplementary Table S2. Reactions were performed in technical duplicate, and melt-curve analysis was used to confirm amplification specificity. Relative expression was calculated using the 2^−ΔΔCt^ method, normalized to 18S rRNA and expressed relative to the indicated control condition.

### Comet assay

DNA damage was assessed using the Comet Assay Kit (3-well slides; Abcam, ab238544) according to the manufacturer’s instructions under alkaline electrophoresis conditions. Briefly, cells were harvested, resuspended in ice-cold PBS at 1×10^6^ cells/mL, mixed with Comet Agarose, and loaded onto the supplied comet slides. Slides were incubated in lysis buffer for 1 h at 4°C, followed by alkaline unwinding for 20 min at 4°C. Electrophoresis was performed in alkaline running buffer at 300 mA for 15 min. Slides were rinsed, fixed in 70% ethanol for 5 min, air-dried, and stained with Vista Green DNA Dye. Images were acquired using a Nikon ECLIPSE Ts2R inverted fluorescence microscope equipped with a 20x objective and controlled using NIS-Elements software (version 5.10; Nikon Instruments). Comet parameters were quantified using the OpenComet v1.3 ImageJ plugin. Tail moment was used as the primary indicator of DNA damage (ref. 39).

### *In silico* analysis of candidate IRE1 RNA endomotifs and XBP1 DNA motif matches

Candidate IRE1 RNase recognition sites were analyzed in human H2AFX (H2A.X), HUWE1, BRCA2, SPARC, and BLOC1S1 mRNA sequences. Reference transcript sequences were obtained from Ensembl using the GRCh38 human genome assembly: H2AFX-202 (ENST00000530167; ENSG00000188486; Ensembl release 106), HUWE1-202 (ENST00000262854; ENSG00000086758; Ensembl release 111), BRCA2-201 (ENST00000380152; ENSG00000139618; Ensembl release 107), SPARC-201 (ENST00000231061; ENSG00000113140; Ensembl release 110), and BLOC1S1-203 (ENST00000548925; ENSG00000135441; Ensembl release 110). Each transcript sequence was screened for the canonical IRE1 RNase endomotif, CUGCAG, and the surrounding sequence context was evaluated for its capacity to form an XBP1-like stem-loop structure.

For each CUGCAG-containing site, a sequence window comprising 40-50 nucleotides upstream and downstream of the motif was extracted and analyzed using the RNAfold web server (ViennaRNA package; accessed July 20, 2026) under default parameters. Predicted minimum-free-energy RNA secondary structures were generated for each sequence window. Candidate IRE1 recognition sites were defined as CUGCAG motifs positioned within the loop region of a predicted stem-loop structure and supported by a flanking paired stem. Predicted structures were visualized and used to map candidate IRE1 endomotifs in H2A.X, HUWE1, BRCA2, SPARC, and BLOC1S1 transcripts. This *in silico* analysis was used to prioritize candidate transcripts for CLIP-qRT-PCR and to evaluate the structural plausibility of IRE1 RNase recognition (**Fig. S4**).

To assess whether the HUWE1 regulatory region contains candidate XBP1 DNA motif matches, the genomic sequence spanning −2,000 to +200 bp relative to the annotated transcription start site of human HUWE1-202 (ENST00000262854; HUWE1 gene ID ENSG00000086758; GRCh38, Ensembl release 116; chromosome X:53,532,096–53,688,752; reverse strand) was retrieved from Benchling. The sequence was analyzed in the 5′→3′ orientation of HUWE1 transcription. Candidate XBP1 motif matches were identified using the JASPAR CORE human XBP1 position-frequency matrix MA0844.2 at a relative-score threshold of 80% (ref. 40). Predicted sites were reported according to their position relative to the annotated TSS, matched sequence, motif orientation, raw score, and relative score (**Fig. S5b**).

### CLIP-RT-qPCR

Cross-linking immunoprecipitation followed by reverse-transcription quantitative PCR (CLIP-RT-qPCR) was performed by Creative BioMart (Shirley, NY, USA) using formaldehyde-fixed MDA-MB-231 cell pellets from non-target control and XBP1-knockdown cells treated with PA and etoposide. IRE1-associated RNAs were immunoprecipitated using an anti-IRE1 antibody, with IgG immunoprecipitation and input lysate included as negative and input controls, respectively. Following proteinase K treatment, reverse cross-linking, and RNA purification, cDNA was analyzed by SYBR Green RT-qPCR for H2A.X, HUWE1, and BRCA2. Primer sequences, CLIP conditions, qPCR reaction composition, and cycling parameters are provided in Supplementary Methods and Table S2.

### Statistical analysis

Statistical analyses were performed using GraphPad Prism version 11 (GraphPad Software, Boston, MA, USA) and Microsoft Excel. Comet-assay data were analyzed using two-sided Mann–Whitney U tests in GraphPad Prism 11. For western blot densitometry, RT-qPCR, and CLIP-qRT-PCR analyses, fold-change or relative-enrichment values were log10-transformed before statistical testing. Two-sided Welch’s t-tests were then performed in Microsoft Excel. Welch’s test was selected because it does not assume equal variances between groups. Data are presented as mean ± SEM unless otherwise stated. All tests were two-sided, and p < 0.05 was considered statistically significant. The number of independent biological replicates used for each experiment are indicated in the corresponding figure legends.

## Results

### IRE1/XBP1 signaling limits DNA damage accumulation during combined PA & etoposide stress

Previously our laboratory and others showed that PA activates the UPR (ref. 35-37). In this study, we investigated the connection between UPR signaling and the DDR, specifically the role of IRE1 in promoting DNA damage repair through the regulation of histone H2A.X. Because XBP1 is a principal downstream effector of IRE1-mediated signaling, we subsequently assessed its contribution to DNA damage repair. We first exposed WT MDA-MB-231 cells to PA (which activates IRE1) (ref.37) and the DNA damage-inducing chemotherapeutic etoposide (ETP). We then used the Comet assay to compare DNA damage accumulation in cells where XBP1 was knocked down (KD) by siRNA to cells receiving a non-targeting siRNA (NT) control. There was a significant increase in DNA damage in XBP1 KD cells as compared with control (**Fig. 1a; green vs. blue**). In contrast, XBP1 KD did not significantly alter DNA damage in WT cells exposed to ETP alone (**Fig. S1a; green vs. blue**), indicating that the DNA-protective contribution of XBP1 is most apparent during combined lipotoxic and genotoxic stress. DNA damage accumulation in IRE1^−/−^ cells was greater than in WT NT control cells under both PA+ETP (**Fig. 1a; yellow/orange vs. blue**) and ETP-only treatment (**Fig. S1a; yellow/orange vs. blue**), regardless of XBP1 KD. In IRE1^−/−^ cells, XBP1 KD did not further increase DNA damage under PA+ETP treatment (**Fig. 1a; yellow versus orange**), indicating that the increase in DNA damage observed following XBP1 depletion in WT cells required intact IRE1. Indeed, re-expression of IRE1 in IRE1^−/−^ cells significantly reduced DNA damage accumulation relative to empty-vector controls under PA+ETP treatment (**Fig. 1b; green vs. blue**). In contrast, under ETP-only treatment, IRE1 re-expression did not significantly reduce DNA damage relative to empty-vector controls (**Fig. S1b; p = 0.0684**). These findings indicate that the DNA-protective effect of IRE1 re-expression is robust under combined PA+ETP stress but was not statistically significant under ETP treatment alone.

**Figure 1:**
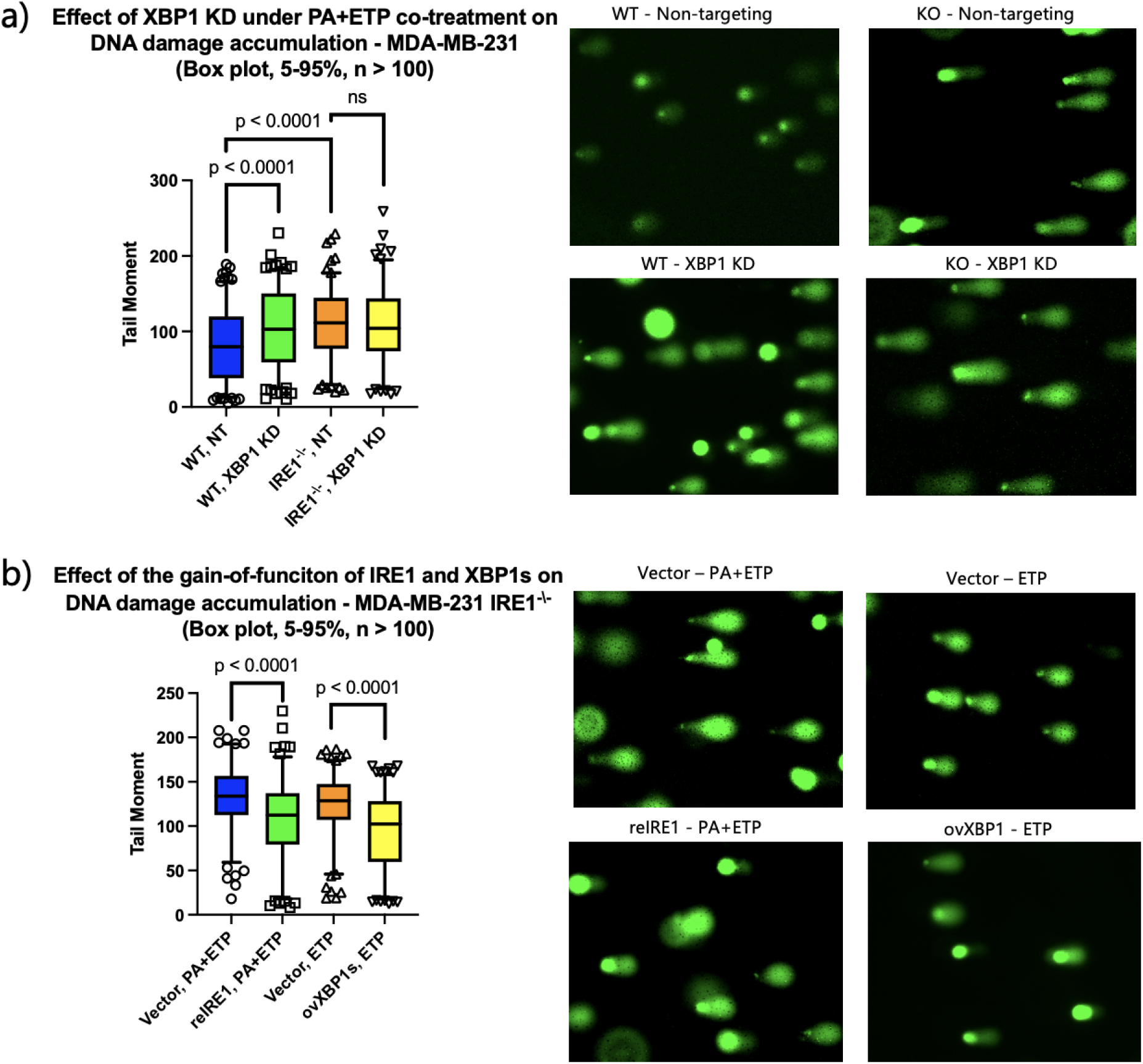
Effects of IRE1 on DNA Damage Accumulation under PA and ETP Exposure: Comet assay of MDA-MB-231 cells. Cells were co-treated with etoposide (ETP, 10 μM) and PA (0.1 mM) as well as ETP only (Fig. 1S). a) DNA damage following XBP1 KD or treatment with a non-targeting siRNA in MDA-MB-231 WT and IRE1^−/−^ cells; b) Re-expression of IRE1 (reIRE1) and overexpression of XBP1s (ovXBP1s) in MDA-MB-231 IRE1^−\−^ cells. Each condition was measured in 3 independent runs with > 50 cells measured per run. Plots show median (horizontal line), 25th and 75th percentiles (bottom and top of the box, respectively), and 5th and 95th percentiles (bottom and top error bars). A representative image for each condition is also shown.

Given that PA activation of IRE1 leads to splicing of XBP1 (XBP1s), we overexpressed XBP1s in IRE1^−/−^ which also reduced DNA damage accumulation relative to empty-vector control under ETP alone treatment (**Fig. 1b; yellow vs. orange**). Next, to assess whether XBP1 depletion altered IRE1 RNase activity, we measured the established RIDD targets BLOC1S1 and SPARC. In WT cells, PA+ETP reduced BLOC1S1 and SPARC mRNA in XBP1 KD cells relative to the corresponding ETP-only condition (**Fig. S2a-b; green vs. blue**), consistent with enhanced IRE1 RNase/RIDD activity under combined stress. This pattern was absent in IRE1^−/−^ cells, in which PA+ETP treatment instead produced a non-significant upward trend in both transcripts (**Fig. S2a-b; yellow vs. orange**), despite similar degrees of XBP1 KD in both WT and IRE1^−/−^(**Fig. S2c-d**). Previous study showed that genotoxic stress, including ETP exposure, can activate RIDD and promote decay of BLOC1S1 and SPARC without inducing XBP1 mRNA splicing in mouse embryonic fibroblasts (ref. 32). Our data extend this observation to MDA-MB-231 cells and suggest that PA co-treatment, together with XBP1 depletion, further favors RIDD-mediated loss of target mRNAs. Thus, reduced XBP1 under combined stress is associated with both increased DNA damage and IRE1-dependent reduction of canonical RIDD-target mRNAs. Consistent with the genetic IRE1-loss phenotype, treatment of WT cells with the IRE1 RNase inhibitor 4μ8C during ETP exposure increased DNA-damage accumulation relative to DMSO control (**Fig. S3**). Together, these genetic and pharmacologic findings support a protective role for IRE1 in limiting DNA-damage accumulation in MDA-MB-231 cells during genotoxic stress, with a more pronounced effect observed during combined PA+ETP exposure.

### IRE1 status modifies the effect of XBP1 depletion on PA-dependent H2A.X and γH2A.X abundance

To connect the UPR and DNA damage response, we next examined the H2A.X regulation, a variant of histone H2A, that is phosphorylated at Ser139 (becoming γH2A.X) in response to DNA DSB. γH2A.X functions both as a marker of DNA damage and as a platform for recruitment and propagation of DNA repair signaling (ref. 16-17). Building on our finding that PA+ETP treatment in IRE1^−/−^ cells with re-expression of IRE1 as well as XBP1s overexpression reduced DNA damage accumulation (**Fig. 1b**), we investigated whether PA-activated IRE1 modulates DNA damage and repair through changes in H2A.X or γH2A.X protein levels.

In IRE1 WT cells, XBP1 KD combined with PA+ETP co-treatment significantly reduced H2A.X protein levels compared with ETP alone (**Fig. 2a; green vs. blue**), consistent with our hypothesis that the IRE1-XBP1 pathway regulates H2A.X expression under genotoxic stress. In contrast, in IRE1^−/−^ cells, XBP1 KD did not alter H2A.X protein levels (**Fig. 2a; yellow vs. orange**), whereas, γH2A.X levels were significantly decreased relative to ETP alone (**Fig. 2b; yellow vs. orange**). These findings suggest the effects of XBP1 depletion on total H2A.X and γH2A.X abundance differed according to IRE1 status.

**Figure 2:**
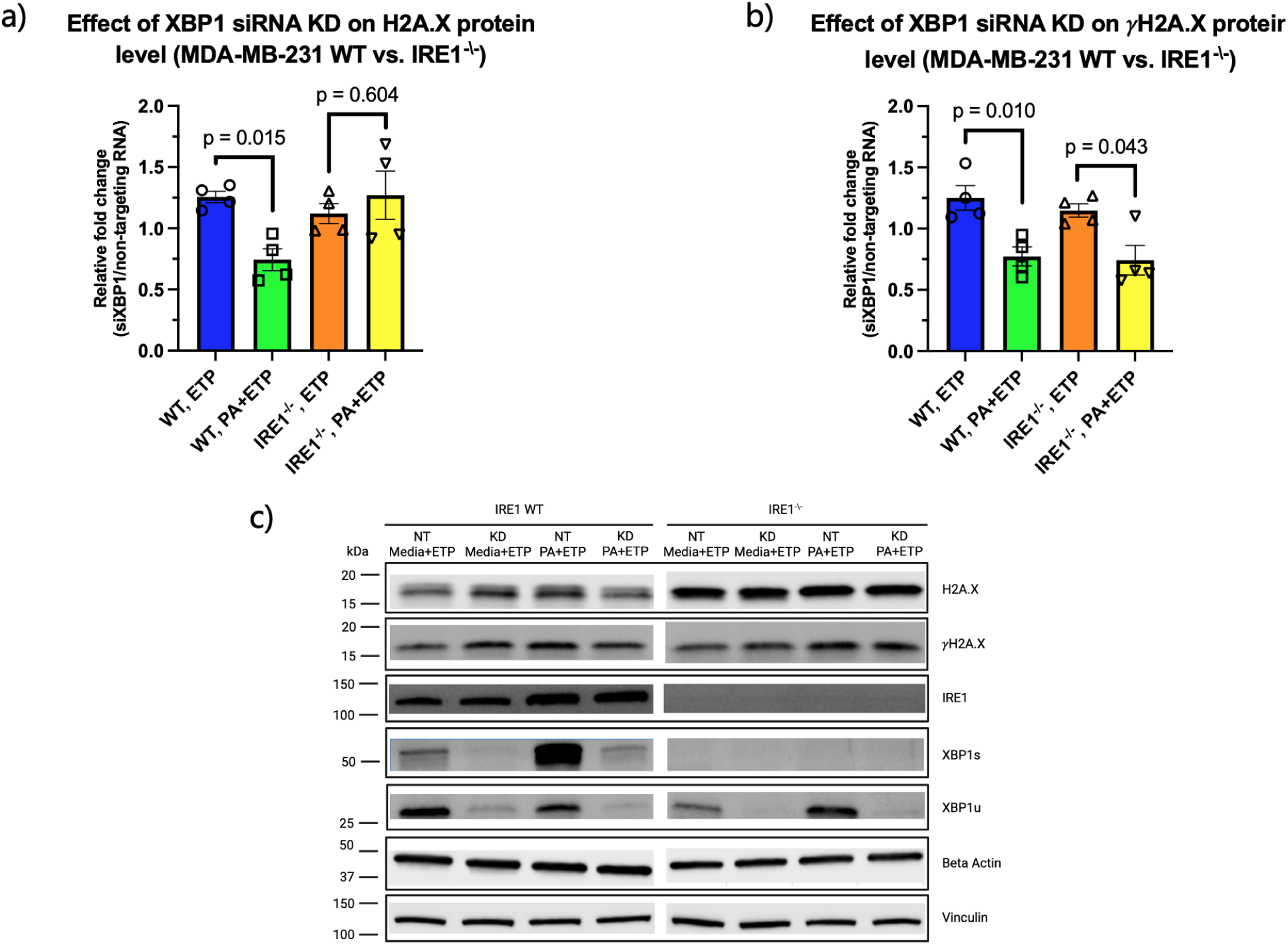
Effects of XBP1 depletion on H2A.X and γH2A.X in IRE1 WT and IRE1^−/−^ cells. a) Quantification of western blots of H2A.X in WT and IRE1^−/−^ cells with XBP1 KD and either co-treatment of PA/ETP or ETP alone. The y-axis is the relative fold change of XBP1 KD to NT control. b) Quantification of western blots of γH2A.X in WT and IRE1^−/−^ cells with XBP1 KD and either co-treatment of PA/ETP or ETP alone. The y-axis is the relative fold change of XBP1 KD to NT control. c) Representative blot for parts a) and b). Data are mean ± SEM from n = 4 independent biological replicates.

### XBP1u, but not XBP1s, alters PA-dependent H2A.X and γH2A.X abundance in IRE1^−/−^ cells

Because XBP1 depletion selectively reduced γH2A.X without altering total H2A.X in IRE1^−/−^ cells, we next investigated whether the individual XBP1 isoforms affected total H2A.X or γH2A.X abundance in the absence of IRE1. We overexpressed XBP1s or unspliced XBP1 (XBP1u) in IRE1^−/−^ cells and compared ETP-only and PA+ETP treatment conditions relative to vector-transfected controls.

XBP1s overexpression did not significantly alter total H2A.X or γH2A.X levels between ETP-only and PA+ETP treatment conditions (**Fig. 3a**). Although γH2A.X levels were modestly higher than those in vector-transfected control cells under both ETP and PA+ETP conditions, these differences were not statistically significant. The γH2A.X/total-H2A.X ratio also did not differ between ETP-only and PA+ETP conditions in XBP1s-overexpressing cells (Source Data Fig. 3a). Thus, XBP1s overexpression was not sufficient to produce a significant PA-dependent change in total H2A.X or γH2A.X abundance in IRE1^−/−^ cells.

**Figure 3:**
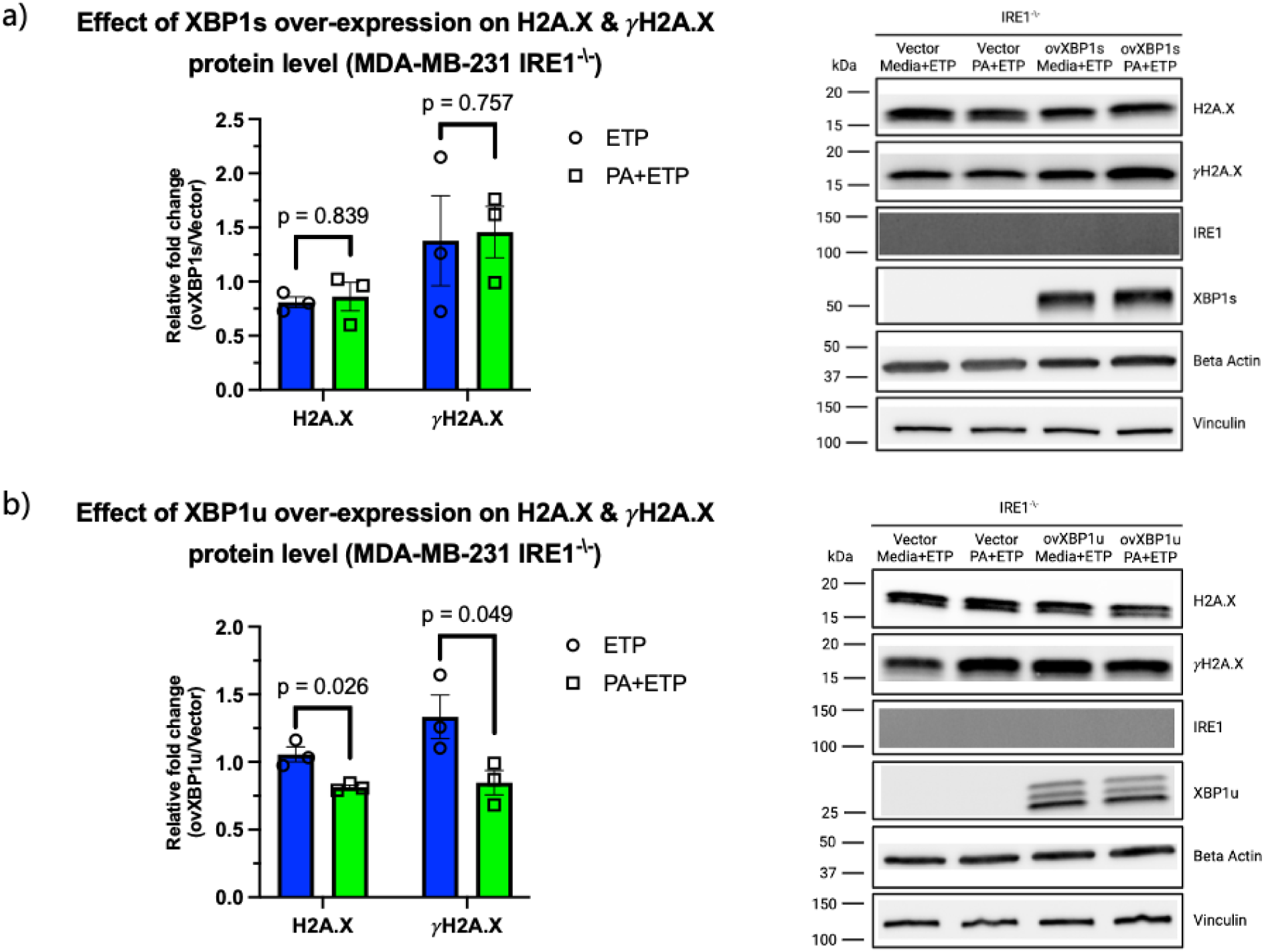
Effects of XBP1s and XBP1u overexpression on H2A.X and γH2A.X in IRE1^−/−^ cells. a) Quantification and representative blot of western blots of H2A.X and γH2A.X in IRE1^−/−^ cells with XBP1s overexpression and either co-treatment of PA/ETP or ETP alone. The y-axis is the relative fold change for cells with XBP1s overexpression relative to empty vector control. b) Quantification and representative blot of western blots of H2A.X and γH2A.X in IRE1^−/−^ cells with XBP1u overexpression and either co-treatment of PA/ETP or ETP alone. The y-axis is the relative fold change for cells with XBP1u overexpression relative to empty vector control. Data are mean ± SEM from n = 3 independent biological replicates.

In contrast, XBP1u overexpression significantly lowers total H2A.X and γH2A.X abundance in PA+ETP-treated cells relative to ETP-only cells in IRE1^−/−^ cells (**Fig. 3b**). The γH2A.X/total-H2A.X ratio did not differ between ETP-only and PA+ETP conditions in XBP1u-overexpressing cells (Source Data Fig. 3b), indicating that the reduction in γH2A.X was proportional to the reduction in total H2A.X. This response differed from the selective reduction in γH2A.X observed following XBP1 depletion in IRE1^−/−^ cells (**Fig. 2b**) and from the lack of a significant PA-dependent response on γH2A.X following XBP1s overexpression (**Fig. 3a**). Together, these findings indicate that XBP1u expression alters the bulk H2A.X/γH2A.X response during PA+ETP exposure in IRE1^−/−^ cells, without a detectable change in the relative phosphorylation state of H2A.X.

### H2A.X as a candidate IRE1-RIDD substrate

To determine whether IRE1 influences H2A.X levels via RIDD, we assessed changes in the H2A.X mRNA abundance. We quantified H2A.X mRNA in WT and IRE1^−/−^ cells under basal conditions (**Fig. 4a**) and following PA+ETP or ETP-only treatment with XBP1 KD or NT control (**Fig. 4b**). At baseline, H2A.X mRNA levels were significantly higher in IRE1^−/−^ cells than in WT, consistent with an IRE1-dependent contribution to H2A.X mRNA regulation. Under PA+ETP treatment, XBP1 KD reduced H2A.X mRNA in WT cells but had no effect in IRE1^−/−^ cells (**Fig. 4b**), indicating that the effect of XBP1 depletion on H2A.X mRNA abundance requires IRE1 and is consistent with a role for IRE1 in regulating H2A.X levels.

**Figure 4:**
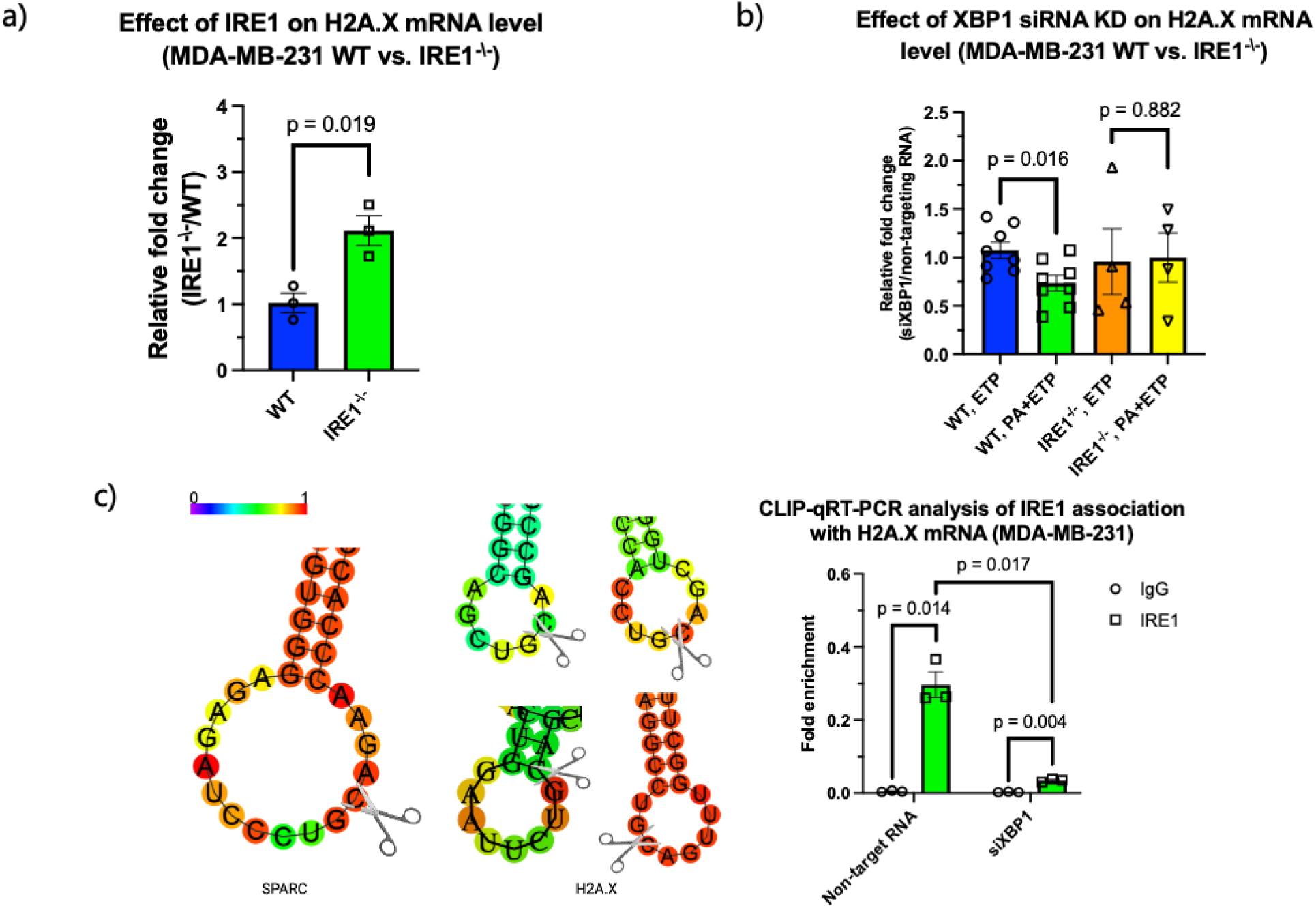
IRE1-H2A.X mRNA Association under PA and ETP Exposure. a) qRT-PCR measuring H2A.X mRNA between WT and IRE1^−/−^ at basal levels. Data are mean ± SEM from n = 3 independent biological replicates. b) qRT-PCR measuring mRNA of H2A.X under PA+ETP co-treatment with XBP1 KD and NT between WT and IRE1^−/−^. Data are mean ± SEM from n = 8 (WT) and n = 4 (IRE1^−/−^) independent biological replicates c) *In silico* RNA secondary-structure predictions for spliced SPARC and H2A.X transcripts generated using RNAfold. SPARC, an established IRE1-RIDD-associated transcript, was included as a positive control. Predicted stem-loop structures containing the putative IRE1 RNase consensus endomotif, CUGCAG, are shown. Scissors indicate predicted IRE1 cleavage sites within the CUGCAG-containing stem-loop structures. Colors indicate base-pairing probabilities from 0 (low probability) to 1 (high probability). CLIP-qRT-PCR analysis of IRE1–H2A.X mRNA association in PA+ETP-treated WT cells with XBP1 KD and NT control. Data are mean ± SEM from n = 3 independent biological replicates.

RIDD-mediated regulation of H2A.X requires physical association between IRE1 and its mRNA target. Using CLIP-qRT-PCR in WT cells treated with PA+ETP and transfected with XBP1 KD or NT control, we observed significant enrichment of H2A.X mRNA in IRE1 immunoprecipitates relative to IgG control (**Fig. 4c**), demonstrating that H2A.X transcripts associate with IRE1 under PA+ETP-treatment. Notably, this H2A.X enrichment was reduced upon XBP1 KD (**Fig. 4c**). Because XBP1 KD also reduces total H2A.X mRNA levels relative to NT control (**Fig. 4b; green**), we normalized CLIP enrichment to input H2A.X mRNA levels in each condition. The reduction in H2A.X association with IRE1 remained significant after input normalization, indicating that XBP1 KD decreases IRE1–H2A.X association independent of the accompanying reduction in total H2A.X transcript abundance. Together, these data identify H2A.X mRNA as an IRE1-associated transcript whose abundance and IRE1 association are influenced by IRE1 status and XBP1 depletion in PA+ETP-treated cells. These findings support H2A.X as a candidate IRE1-regulated transcript and possible RIDD target. Future experiments are needed to test whether H2A.X is directly cleaved by IRE1 through RIDD.

### IRE1 engages additional DNA damage response-related mRNA: HUWE1 and BRCA2 identified by CLIP-qRT-PCR

Having established that H2A.X mRNA associates with IRE1 under PA + ETP-treatment condition and in XBP1-dependent manner (**Fig. 4**), we next asked whether IRE1 also engages additional DDR-related mRNAs mechanistically linked to H2A.X/γH2A.X. We prioritized HUWE1 and BRCA2 because both are established regulators of H2A.X-associated DNA damage signaling. HUWE1 is a HECT-domain E3 ubiquitin ligase that monoubiquitinates H2A.X, thereby supporting γH2A.X signaling, recruitment of repair factors, and restart of stalled replication forks. HUWE1 is recruited to stalled replication forks through PCNA, where it monoubiquitinates H2A.X and supports replication-stress signaling and fork recovery (ref. 18). BRCA2 functions in homologous recombination and contributes to stabilization of stalled replication forks (ref. 19, 38).

At baseline, BRCA2 mRNA levels were significantly increased in IRE1^−/−^ cells relative to WT cells, paralleling the increase observed for H2A.X mRNA. In contrast, HUWE1 mRNA levels were significantly decreased in IRE1^−/−^ cells relative to WT cells (**Fig. 5a**). These divergent expression patterns indicate that loss of IRE1 differentially affects basal abundance of DDR-associated transcripts: the elevated BRCA2 and H2A.X mRNA levels are consistent with an IRE1-dependent contribution to their transcript regulation, whereas the reduced HUWE1 mRNA level is inconsistent with a simple IRE1-mediated decay model and may reflect indirect transcriptional or post-transcriptional regulation.

**Figure 5:**
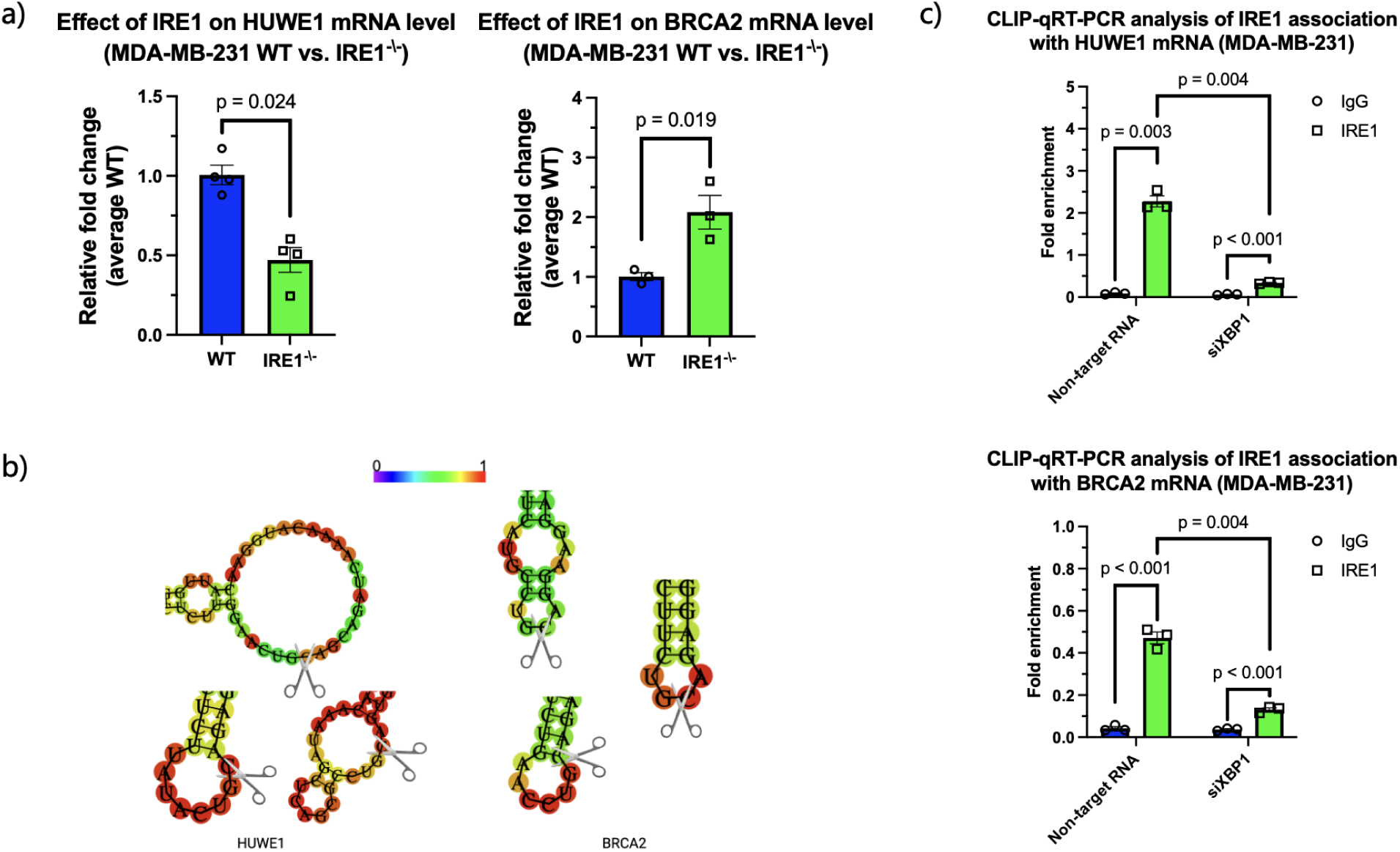
IRE1-HUWE1 and BRCA2 mRNA Association under PA and ETP Exposure. a) qRT-PCR measuring HUWE1 and BRCA2 mRNA between WT and IRE1^−/−^ at basal levels. b) *In silico* RNA secondary-structure predictions for spliced SPARC, HUWE1, and BRCA2 transcripts generated using RNAfold. SPARC, an established IRE1-RIDD-associated transcript, was included as a positive control. Predicted stem-loop structures containing the putative IRE1 RNase consensus endomotif, CUGCAG, are shown. Scissors indicate predicted IRE1 cleavage sites within the CUGCAG-containing stem-loop structures. Colors indicate base-pairing probabilities from 0 (low probability) to 1 (high probability). c) CLIP-qRT-PCR analysis of IRE1–HUWE1 and IRE1-BRCA2 mRNA association in PA+ETP-treated WT cells with XBP1 KD and NT control. Data are mean ± SEM from n = 3 independent biological replicates.

*In silico* analysis of spliced HUWE1 and BRCA2 mRNAs revealed multiple CUGCAG-containing stem-loop structures matching the reported IRE1 RNase endomotif (**Fig. 5b**), providing a rationale for including these transcripts in our CLIP panel to test whether IRE1 associates with core DNA-repair mRNAs beyond H2A.X. However, the presence of candidate motifs alone does not establish direct IRE1 cleavage. In IRE1 WT cells, CLIP-qRT-PCR revealed significant enrichment of both HUWE1 and BRCA2 mRNAs in IRE1 immunoprecipitates relative to IgG controls (**Fig. 5c**), demonstrating that these DDR-related transcripts associate with IRE1 under PA + ETP treatment. The extent of HUWE1 and BRCA2 enrichment was comparable to that of H2A.X mRNA (**Fig. 4c**), supporting that IRE1 associates with multiple DDR-related transcripts under combined ER and genotoxic stress. Notably, HUWE1 and BRCA2 enrichment in IRE1 immunoprecipitates was reduced upon XBP1 KD (**Fig. 5c**), suggesting that these RNA interactions are modulated by XBP1 status under PA+ETP treatment.

Together, these findings identify HUWE1 and BRCA2 as IRE1-associated DDR transcripts with distinct basal expression phenotypes in IRE1^−/−^ cells. BRCA2, like H2A.X, is a candidate for IRE1-dependent RNA regulation, whereas HUWE1 showed reduced basal abundance and a treatment-dependent relative response to XBP1s overexpression in IRE1^−/−^ cells (**Fig. S5**). Further experiments are required to determine whether either transcript undergoes direct IRE1 RNase-mediated cleavage.

## Discussion

Our findings identify PA-activated IRE1 as a context-dependent regulator of the DNA damage response in MDA-MB-231 cells. The protective effects of IRE1 re-expression and XBP1s, together with the sensitization caused by XBP1 depletion specifically during combined PA+ETP exposure, support an adaptive role for the IRE1–XBP1 axis under combined lipotoxic and genotoxic stress. This context dependence is important because XBP1 depletion did not measurably affect DNA damage under ETP alone. Although IRE1 re-expression significantly reduced DNA damage under PA+ETP treatment, its effect under ETP-only treatment did not reach statistical significance. Thus, our data support a robust IRE1-dependent protective response during combined lipotoxic and genotoxic stress, while XBP1-dependent protection was most apparent in the PA-associated stress context.

XBP1 depletion was also associated with IRE1-dependent reduction of the established RIDD-associated transcripts BLOC1S1 and SPARC under PA+ETP treatment. This finding is consistent with altered partitioning of IRE1 RNase output between XBP1 mRNA processing and RIDD-associated RNA decay. Dufey et al. similarly demonstrated that genotoxic stress can promote IRE1-dependent decay of BLOC1S1 and SPARC without robust XBP1 splicing in mouse embryonic fibroblasts (ref. 32). However, the present steady-state mRNA measurements do not establish whether RIDD is directly protective, detrimental, or simply concurrent with the DNA-damage phenotype. Therefore, the relationship between IRE1 RNase output and DNA-damage resolution is likely more complex than a binary XBP1s-versus-RIDD model.

The H2A.X/γH2A.X data indicate that XBP1-dependent responses to PA+ETP differ according to IRE1 status. In WT cells, XBP1 depletion reduced total H2A.X under PA+ETP relative to ETP alone. In IRE1^−/−^ cells, XBP1 depletion did not alter total H2A.X but reduced γH2A.X, indicating that the H2A.X-associated response to XBP1 depletion differs in the presence and absence of IRE1. XBP1 isoform overexpression was examined specifically in IRE1^−/−^ cells. Neither XBP1s nor XBP1u altered the γH2A.X/total-H2A.X ratio between ETP-only and PA+ETP conditions. However, XBP1u reduced both total H2A.X and γH2A.X under PA+ETP, consistent with a proportional change in bulk H2A.X abundance rather than selective modulation of H2A.X phosphorylation. In contrast, XBP1s did not significantly alter bulk H2A.X or γH2A.X between treatment conditions but reduced DNA-damage accumulation, suggesting that its protective effect may involve other DNA-damage-response or stress-adaptation programs.

Our RNA analyses identify H2A.X as an IRE1-associated and IRE1-sensitive candidate DDR transcript. The elevated basal abundance of H2A.X mRNA in IRE1^−/−^ cells, the IRE1-dependent reduction in H2A.X mRNA following XBP1 depletion under PA+ETP treatment, and the enrichment of H2A.X mRNA in IRE1 immunoprecipitates together support a role for IRE1 in regulating H2A.X mRNA. Importantly, after normalizing the H2A.X CLIP signal to H2A.X mRNA abundance in the corresponding input lysates, IRE1-associated H2A.X mRNA remained reduced following XBP1 depletion. This result indicates that the reduced H2A.X signal recovered in the IRE1 immunoprecipitate after XBP1 depletion was not explained solely by lower input H2A.X mRNA abundance, but was also consistent with reduced relative enrichment of H2A.X mRNA in the IRE1 immunoprecipitate. Because IRE1 can mediate RIDD of selected RNAs, the increased basal H2A.X mRNA abundance in IRE1^−/−^ cells, the predicted CUGCAG-containing stem-loop motif, and H2A.X association with IRE1 raise the possibility that H2A.X may be a RIDD substrate. However, these observations do not demonstrate direct IRE1 RNase-mediated cleavage or RIDD-dependent turnover. H2A.X should therefore be considered an IRE1-associated candidate RIDD substrate, pending direct cleavage and RNA-decay experiments.

The identification of HUWE1 and BRCA2 as additional IRE1-associated transcripts suggests that IRE1 may engage a broader network of H2A.X-linked DDR mRNAs under combined lipotoxic and genotoxic stress. Their divergent basal expression patterns after IRE1 loss indicate that IRE1 effects on DDR-associated RNAs are transcript specific and cannot be explained by a uniform RNA-decay mechanism. Whereas BRCA2, like H2A.X, showed increased basal mRNA abundance following IRE1 loss, HUWE1 showed reduced basal abundance and a treatment-dependent relative response to XBP1s overexpression in IRE1^−/−^ cells. The predicted CUGCAG-containing stem-loop motifs and IRE1 association provide a rationale for further studies to determine whether HUWE1 or BRCA2 undergoes direct IRE1 RNase cleavage or RIDD-mediated turnover.

Collectively, these data support a model in which PA-activated IRE1 contributes to genome-protective responses through XBP1-dependent signaling and potentially through RNase-associated regulation of DDR-related transcripts, with associated changes in bulk H2A.X/γH2A.X dynamics. Future studies should directly measure H2A.X, HUWE1, and BRCA2 mRNA decay kinetics and IRE1-dependent cleavage; use genetic separation-of-function mutants to distinguish XBP1 splicing from RIDD; and determine whether these mechanisms influence DNA repair capacity and response to genotoxic therapy across additional TNBC models and other tumor types.

## Declaration

### Ethical Approval

Not applicable.

### Consent to Publish declaration

Not applicable.

## Author Contributions

All authors contributed to methodology, conceptualization, data acquisition, investigation and manuscript reviewing. Kevin Chen wrote the original draft.

## Data Sharing

The data that support the findings of this study are available from the corresponding author upon reasonable request.

