## Supplemental File for "Palmitate-activated IRE1-XBP1 signaling limits DNA damage and is associated with H2A.X-linked DNA damage response transcripts in triple-negative breast cancer cells"

### Supplementary Information

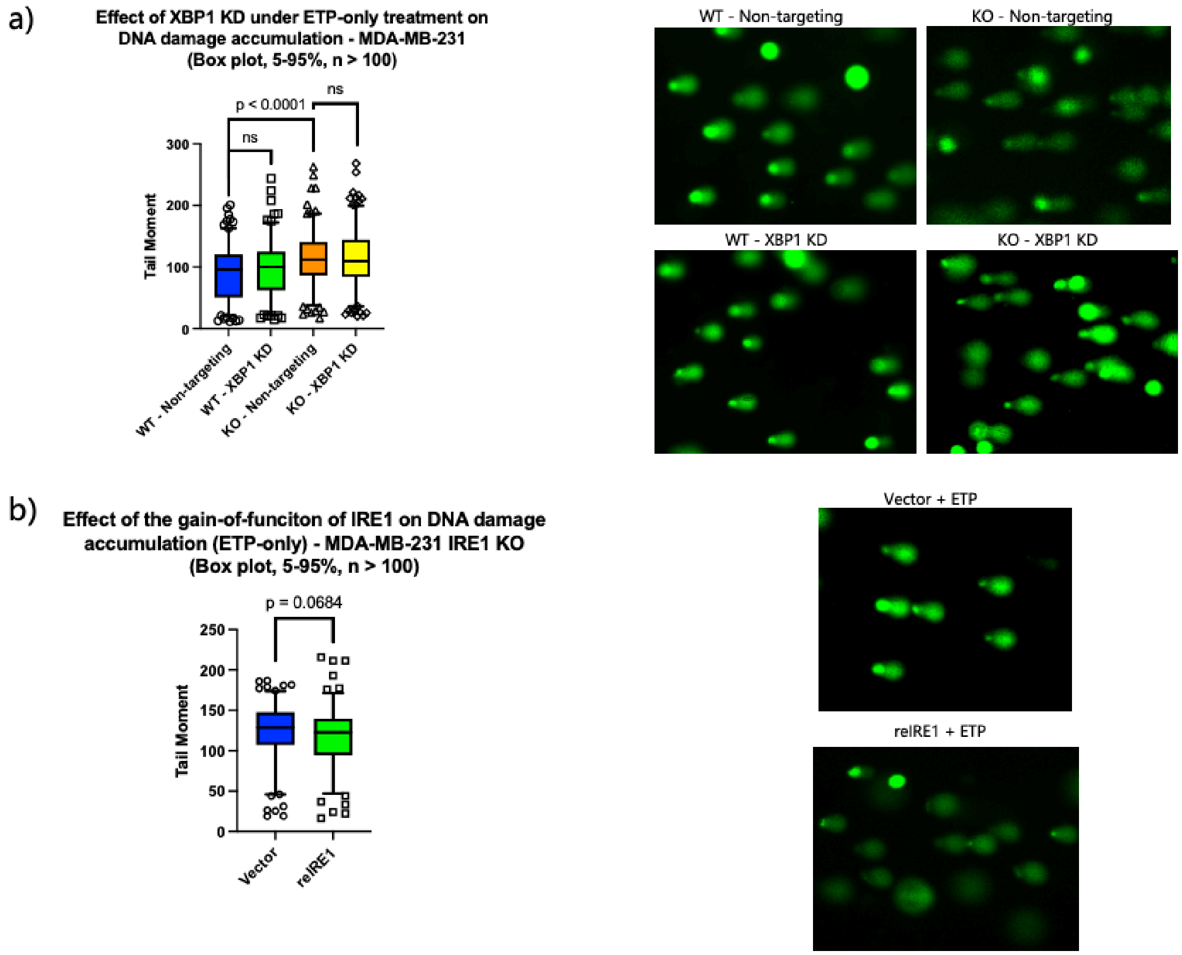


**Figure S1. Effects of ETP alone on DNA damage in MDA-MB-231 WT and IRE1$^{-/-}$ cells.** MDA-MB-231 WT and IRE1$^{-/-}$ cells were treated with etoposide (ETP, 10 μM). (a) DNA damage was assessed following transfection with XBP1-targeting siRNA (XBP1 KD) or non-targeting siRNA (NT) in WT and IRE1$^{-/-}$ cells. (b) DNA damage was assessed following re-expression of IRE1 in MDA-MB-231 IRE1$^{-/-}$ cells. Each condition was measured in three independent experiments, with > 50 cells quantified per experiment. Plots show the median (horizontal line), 25th and 75th percentiles (bottom and top of the box, respectively), and 5th and 95th percentiles (error bars). Representative comet images are shown for each condition. XBP1 knockdown did not significantly alter DNA damage in WT cells under ETP-only treatment. In contrast, re-expression of IRE1 did not significantly reduce ETP-induced DNA damage in IRE1$^{-/-}$ cells ($p = 0.0684$). This result contrasts with the significant reduction in DNA damage observed under combined PA+ETP treatment ($P < 0.0001$; **Fig. 1b**).

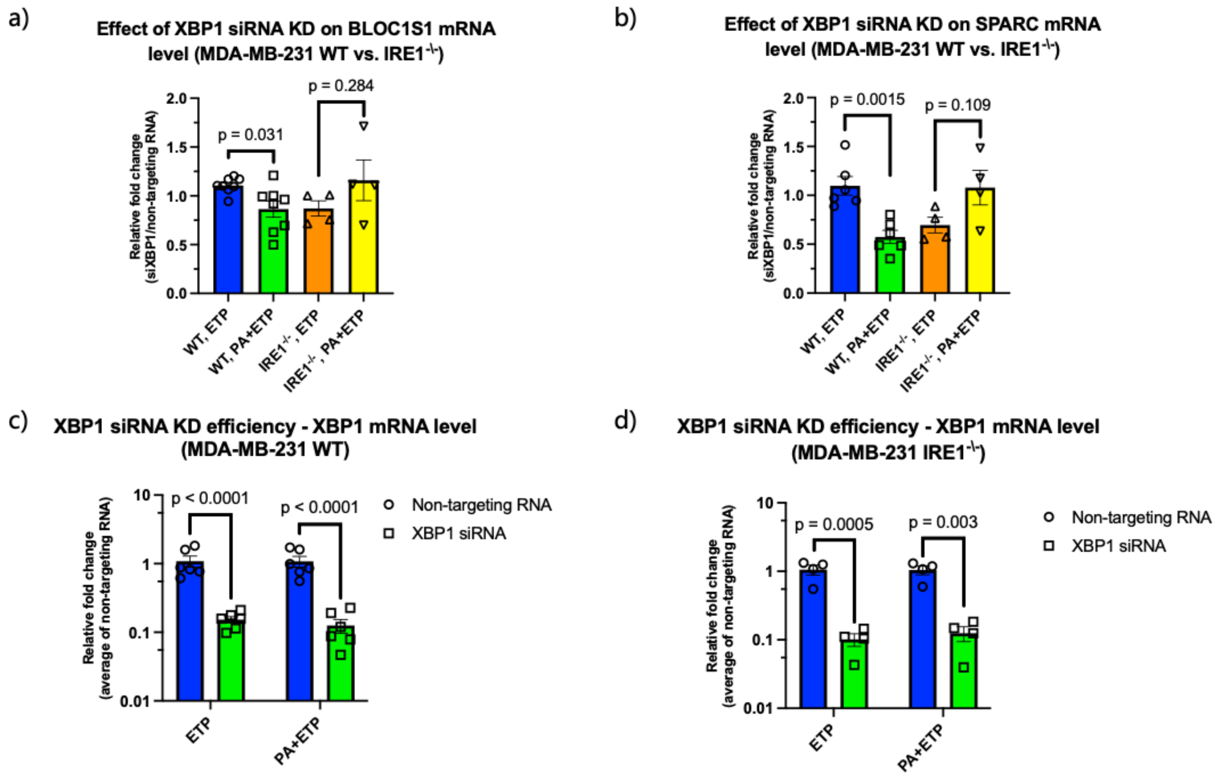


**Figure S2. XBP1 depletion shifts RIDD-target expression in an IRE1-dependent manner.** MDA-MB-231 WT and $IRE1^{-/-}$ cells were transfected with XBP1 KD or NT siRNA and treated with ETP alone or in combination with PA. qRT-PCR was used to measure the relative mRNA expression of the established RIDD targets (a) BLOC1S1 and (b) SPARC, as well as XBP1 mRNA in (c) WT and (d) $IRE1^{-/-}$ cells. Data are presented as the XBP1 KD/NT ratio for each treatment condition for (a) and (b), normalized to average NT in (c) and (d). The y-axis in (c) and (d) is displayed on a base-10 logarithmic scale to accommodate the dynamic range of XBP1 expression. Data are mean ± SEM. For IRE1 WT cells, n = 8 for *BLOC1S1* and n = 6 for *SPARC* and *XBP1*. For $IRE1^{-/-}$ cells, n = 4 for all transcripts. In WT cells, PA+ETP reduced BLOC1S1 and SPARC mRNA relative to ETP alone, consistent with increased IRE1 RNase/RIDD activity under combined lipotoxic and genotoxic stress. In $IRE1^{-/-}$ cells, PA+ETP produced an opposite, non-significant upward trend in BLOC1S1 and SPARC expression, supporting the IRE1 dependence of the PA+ETP-associated reduction observed in WT cells. XBP1 mRNA was significantly reduced by XBP1 KD in both WT and $IRE1^{-/-}$ cells under ETP and PA+ETP conditions, confirming knockdown efficiency.

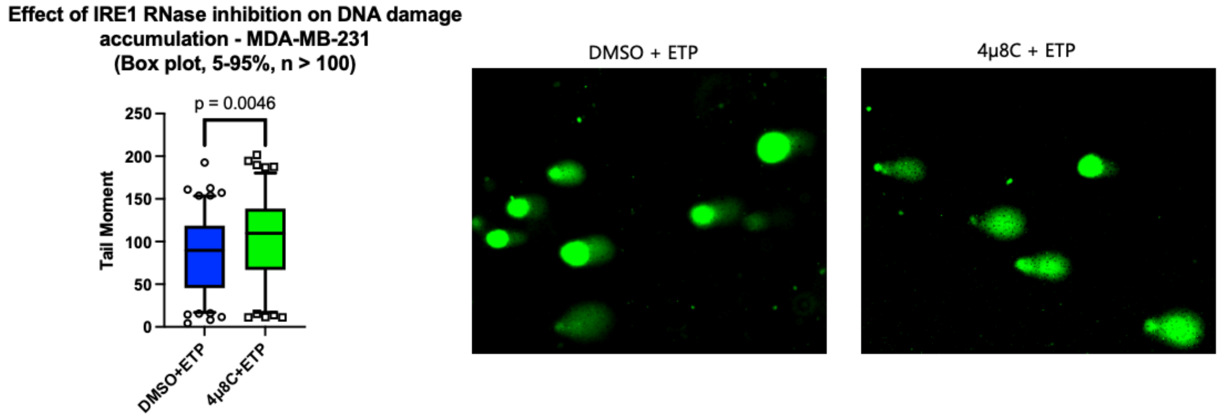


**Figure S3. Pharmacologic inhibition of IRE1 RNase activity increases ETP-associated DNA-damage accumulation in MDA-MB-231 WT cells.** To pharmacologically assess the contribution of IRE1 RNase activity during genotoxic stress, MDA-MB-231 WT cells were treated with 4μ8C or DMSO during ETP exposure. Each condition was measured in 3 independent runs with > 50 cells measured per run. Plots show median (horizontal line), 25th and 75th percentiles (bottom and top of the box, respectively), and 5th and 95th percentiles (bottom and top error bars). A representative image for each condition is also shown. Relative to DMSO+ETP controls, 4μ8C+ETP treatment significantly increased DNA-damage accumulation in the Comet assay. These results provide pharmacologic support for the genetic evidence that IRE1 contributes to limiting DNA-damage accumulation during ETP exposure.

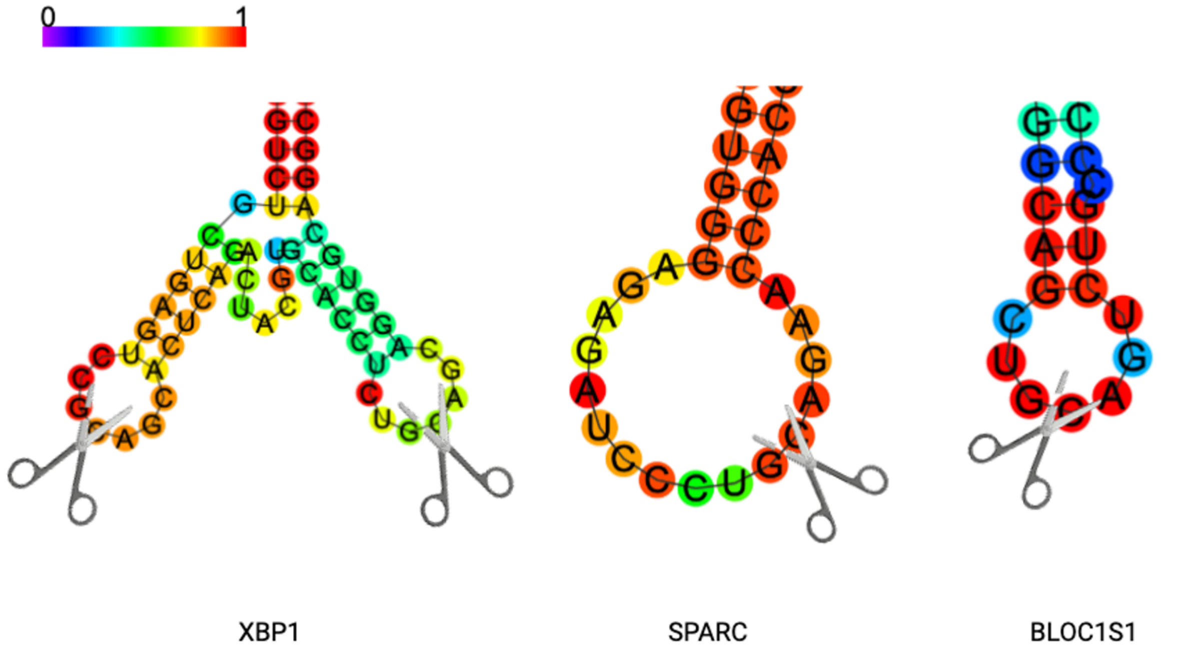

**Figure S4. Predicted IRE1 RNase endomotifs and local RNA secondary structures in established IRE1/RIDD-associated transcripts.** RNA sequences surrounding CUGCAG motifs in human XBP1, SPARC, and BLOC1S1 transcripts were analyzed using RNAfold (ViennaRNA package). Sequence windows extending approximately 40–50 nucleotides upstream and downstream of each CUGCAG motif were folded using default parameters. Predicted minimum-free-energy structures are shown, with the CUGCAG motif highlighted. Colors indicate base-pairing probabilities from 0 (low probability) to 1 (high probability). Candidate IRE1 RNase recognition sites were defined as CUGCAG motifs positioned in the loop of a predicted stem-loop structure with a flanking paired stem. These computational predictions provide structural context for the known IRE1-associated transcripts but do not establish direct IRE1 cleavage or RIDD-mediated decay.

a)

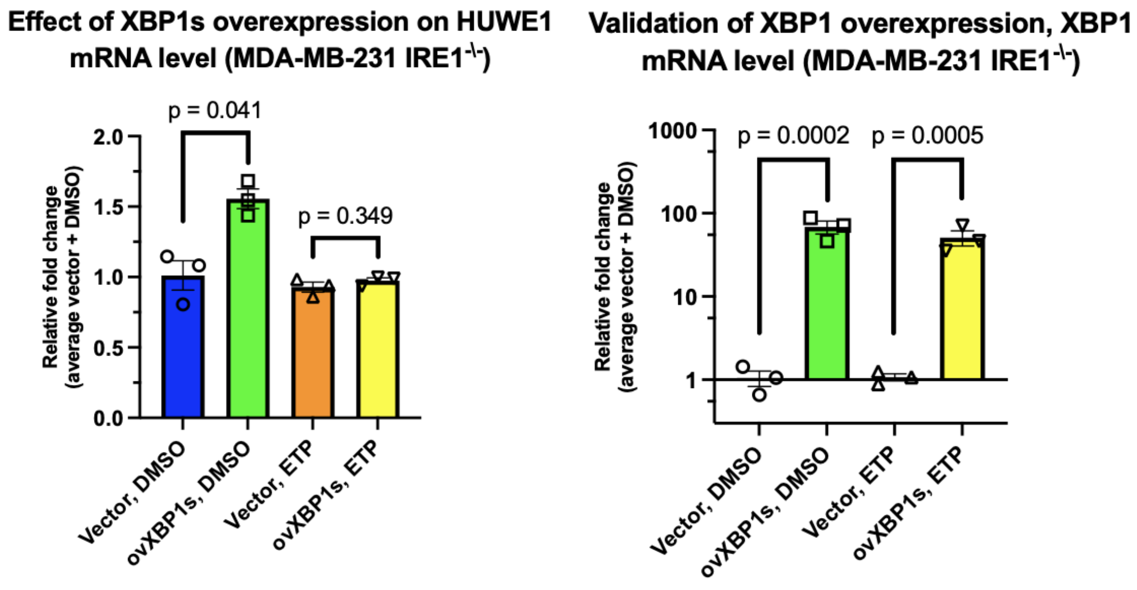


b)

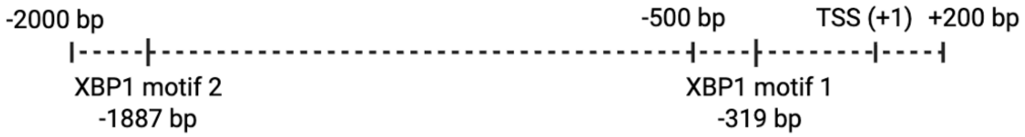


| Candidate site | Predicted sequence | Relative to TSS | Strand | Relative score | JASPAR matrix |
|---|---|---|---|---|---|
| XBP1 motif 1 | 5'-CCCACGTCGTC-3' | −319 to −309 bp | + | 85.76% | MA0844.2 |
| XBP1 motif 2 | 5'-GATACCTCATC-3' | −1887 to −1877 bp | + | 81.49% | MA0844.2 |

**Figure S5. XBP1s overexpression validation, HUWE1 mRNA response, and predicted XBP1 motif matches in the HUWE1 TSS-proximal regulatory region.** (a) MDA-MB-231 IRE1$^{-\backslash-}$ cells transfected with XBP1s-expression plasmid or vector control were treated with DMSO or ETP. Total XBP1 and HUWE1 mRNA were measured by RT-qPCR and normalized to 18S rRNA. HUWE1 values are shown as fold change relative to matched vector control plus DMSO. The vector-normalized HUWE1 response to XBP1s overexpression was greater with DMSO than ETP treatment. (b) JASPAR scanning of the −2,000 to +200 bp interval relative to the TSS of the selected HUWE1 transcript identified two candidate XBP1 motif matches using matrix MA0844.2 at a relative-score threshold of 80%. Motif locations, sequences, orientations, and scores are shown. Predicted motifs do not establish XBP1s occupancy or direct transcriptional regulation.

| Target | Host/clonality | Vandor | Catalog no. | Expected molecular weight (kDa) |
|---|---|---|---|---|
| Vinculin | Rabbit mAb | Cell Signaling Technology | 13901 | 124 |
| β-Actin | Rabbit mAb | Cell Signaling Technology | 8457 | 45 |
| IRE1 | Rabbit mAb | Cell Signaling Technology | 3294 | 130 |
| XBP1 | Rabbit mAb | Abcam | ab220783 | XBP1u ~29 ; XBP1s ~54 |
| H2A.X | Rabbit mAb | Abcam | ab229914 | ~15 |
| γH2A.X | Rabbit mAb | Abcam | ab81299 | ~15 |
| Secondary | Goat polyclonal | Thermo Fisher Scientific | 31460 | - |

**Supplementary Table S1. Primary and secondary antibodies used for western blotting.** The table lists the target antigen, host species and clonality, vendor, catalog number, and expected molecular weight for each antibody used for immunoblot analysis. Antibody working dilutions are described in the Western blotting Methods section.

| Target | Forward (5'-3') | Reverse (5'-3') |
|---|---|---|
| H2A.X | GGCCTCCAGTTCCCAGTG | TCAGCGGTGAGGTACTCCAG |
| HUWE1 | CCAGAAGTTCTTCTTGAGGGTAC | GCCTAAACCGGAGGAACC |
| BRCA2 | TGAAATTAAACGGAAGTTTGC | GAATAAAAGCCCCTAAACCC |
| XBP1 | TTACGAGAGAAAACTCATGGCC | GGGTCCAAGTTGTCCAGAATGC |
| SPARC | AGCACCCCATTGACGGGTA | GGTCACAGGTCTCGAAAAAGC |
| BLOC1S1 | AGAACTGGGCTCGGAGCAT | AGCTGCCCTTTGTAGACATAT |
| 18S | ATGGCCGTTCTTAGTTGGTG | CGCTGAGCCAGTCAGTGTAG |

**Table S2. Primer sequences used for RT-qPCR and CLIP-qRT-PCR.** Gene-specific forward and reverse primer sequences used to quantify H2A.X, HUWE1, BRCA2, XBP1, SPARC, BLOC1S1, and the reference gene 18S rRNA are listed in the 5′→3′ direction. H2A.X, HUWE1, and BRCA2 primer pairs were used for both RT-qPCR and CLIP-qRT-PCR analyses.

**Supplementary Methods**

Detailed cell-treatment and transfection procedures

MDA-MB-231 cells were seeded one day before transfection in 70-mm tissue-culture-treated dishes (GenClone; cat. no. 25-201) at $8\times10^5$ cells per dish for siRNA experiments or $1\times10^6$ cells per dish for plasmid-transfection experiments. Cells were transfected with 10 nM XBP1-targeting siRNA or non-targeting control siRNA using Lipofectamine 3000 Transfection Reagent (Thermo Fisher Scientific; cat. no. L3000150) according to the manufacturer's instructions. siRNA-transfected cells were incubated for 48 h before chemical treatment.

For siRNA-mediated *XBP1* knockdown, cells were transfected with 10 nM ON-TARGETplus Human *XBP1* siRNA SMARTpool (Horizon Discovery, Cambridge, UK; cat. no. L-009552-00-0005) or 10 nM ON-TARGETplus Non-targeting Control Pool (Horizon Discovery; cat. no. D-001810-10-05) using Lipofectamine 3000 Transfection Reagent (Thermo Fisher Scientific, Waltham, MA, USA; cat. no. L3000150) according to the manufacturer's instructions. Cells were incubated for 48 h after transfection before PA and/or ETP treatment.

For plasmid overexpression and IRE1 re-expression experiments, MDA-MB-231 IRE1$^{-\backslash-}$ cells were transfected with 1,000 ng plasmid DNA per 70-mm dish using Lipofectamine 3000 according to the manufacturer's instructions. Cells received a GenScript custom empty pcDNA3.1(+) vector (SC1317) or GenScript pcDNA3.1+/C-(K)-DYK expression plasmids encoding human IRE1/ERN1 (Clone ID OHu25357; RefSeq NM_001433.5), XBP1s (Clone ID OHu25513; RefSeq NM_001079539.1), or XBP1u (Clone ID OHu26371; RefSeq NM_005080.3). The expression constructs contained C-terminal DYKDDDDK tags. Equal total amounts of plasmid DNA were used across conditions. Cells were incubated for 24 h after transfection before PA and/or ETP treatment.

Sodium palmitate (PA; Sigma-Aldrich, St. Louis, MO, USA; cat. no. P9767) was dissolved in water to prepare a 30 mM stock solution by heating until fully dissolved. Immediately before use, the PA stock was rapidly diluted into complete glucose-supplemented DMEM to a final concentration of 0.1 mM. For PA+ETP experiments, cells were pretreated with 0.1 mM PA for 8 h and then co-treated with 0.1 mM PA and 10 µM ETP (Sigma-Aldrich, St. Louis, MO, USA; cat. no. E1383) for an additional 16 h. ETP-only conditions received 10 µM ETP for 16 h in the absence of PA. Cells not treated with PA received complete culture medium without PA.

For IRE1 RNase-inhibition experiments, WT MDA-MB-231 cells were treated with 30 µM 4µ8C (Sigma-Aldrich, St. Louis, MO, USA; cat. no. SML0949) or volume-matched DMSO vehicle for 3 h before the addition of 10 µM ETP. Cells remained in 4µ8C or DMSO during a subsequent 3-h ETP exposure before processing for Comet assay.

CLIP–RT-qPCR

Formaldehyde-fixed MDA-MB-231 cell pellets were prepared using 0.35% formaldehyde for 10 min at room temperature. Fixation was quenched by addition of 2.5 M glycine at a 1:40 ratio for 5 min at room temperature. Cells were washed with ice-cold PBS, pelleted, snap-frozen in liquid nitrogen, and stored at −80 °C until analysis.

CLIP–RT-qPCR was performed by Creative BioMart (Shirley, NY, USA). Cell pellets were washed twice with ice-cold PBS and lysed in CLIP lysis buffer supplemented with PMSF, protease inhibitor cocktail, low-RNase solution, and DNase. Lysates were incubated on ice for 2 min, sonicated for five cycles of 30 s with 30-s intervals, and centrifuged at 12,000 rpm for 10 min at 4 °C. An aliquot of lysate was retained as the input control.

Clarified lysates were incubated overnight at 4 °C with rotation with anti-IRE1 antibody or control IgG. Immune complexes were captured using magnetic beads for 4 h at 4 °C and washed three times with CLIP binding buffer. Complexes were eluted at 37 °C for 5 min with agitation at 1,000 rpm. Eluates were treated with proteinase K at 56 °C for 15 min followed by 72 °C for 15 min. RNA was extracted with TRIzol and purified using a micro-RNA extraction kit.

qPCR was performed in 20-µL reactions containing 10 µL Power SYBR Green Master Mix, 0.5 µL each of forward and reverse primers (10 µM), 1 µL cDNA, and 8 µL nuclease-free water. Reactions were run with an initial denaturation at 95 °C for 1 min, followed by 40 cycles of 95 °C for 15 s and 63 °C for 25 s, with fluorescence acquisition during each cycle. Melt-curve analysis was performed from 55 °C to 95 °C. Primer pairs targeted HUWE1, H2A.X, and BRCA2; their sequences are provided in Table S1.
